# Identification of intestinal mediators of *Caenorhabditis elegans* DBL-1/BMP immune signaling shaping gut microbiome composition

**DOI:** 10.1101/2023.10.26.564221

**Authors:** Kenneth Trang, Barbara Pees, Siavash Karimzadegan, Rahul Bodkhe, Sabrina Hammond, Michael Shapira

**Author notes:** Present Address: Department of Evolutionary Ecology and Genetics, Zoological Institute, Christian-Albrecht University, Kiel, Germany (Barbara Pees). Present Address: Department of Biological Sciences, Contra Costa College, San Pablo, California, USA. These authors contributed equally to the manuscript: KT, BP, and SK all performed experiments and analyzed data vital to the manuscript, with KT leading with manuscript writing, BP involved in the initial project conception, and SK involved in figure preparation. The order of these three lead-co-authors was determined by slight differences in the extent of contributions.

## Abstract

The composition of the gut microbiome is determined by a complex interplay of diet, host genetics, microbe-microbe competition, abiotic factors, and stochasticity. Previous studies have demonstrated the importance of host genetics in community assembly of the *Caenorhabditis elegans* gut microbiome and identified a pivotal role for DBL-1/BMP immune signaling in determining the abundance of gut *Enterobacteriaceae*, in particular of the genus *Enterobacter*. However, the effects of DBL-1 signaling on gut bacteria were found to depend on its activation in extra-intestinal tissues, suggesting that yet unidentified intestinal factors must mediate these effects. In the present study, we used RNA-seq gene expression analysis of wildtype, *dbl-1* and *sma-3* mutants, and *dbl-1* over-expressors to identify genes regulated by DBL-1/BMP signaling that take part in interactions with gut commensals. Following confirmation of several putative targets by qRT-PCR, we carried out colonization experiments with respective mutants raised on monocultures as well as on defined bacterial communities. These experiments identified five intestinal DBL-1/BMP targets, predicted to be secreted, that showed increased *Enterobacteriaceae* abundance compared to wildtype. The extent of increases was for the most part lower than those seen in DBL-1 pathway mutants, suggesting that identified mediators are components of a DBL-1-regulated antibacterial cocktail, which may additively contribute to shaping of gut microbiome composition.

**IMPORTANCE:** Compared to the roles of diet, environmental availability, or lifestyle in determining gut microbiome composition, that of genetic factors is the least understood and often underestimated. The identification of intestinal mediators acting downstream of DBL-1/BMP signaling to control enteric bacteria, describes a cocktail of effectors with distinct molecular functions, thus offering a glimpse into the genetic logic of microbiome control as well as a list of targets for future exploration of this logic.

## INTRODUCTION

Animals harbor large gut microbial communities (microbiomes) that play important roles in host health and fitness. The composition of these communities is shaped by various factors, including environmental microbial availability, diet, lifestyle, and host genetics (1). In recent years, a greater appreciation is emerging of the roles that host genetics play in the interactions between animals and microbes (2), but overall, host genetics remains less characterized than other determinants of gut microbiome composition. In humans, genome-wide association studies have revealed associations between gene variants and gut microbiome composition, including between variants of the LCT lactase gene and *Bifidobacteriaceae*, thought to be linked through lactose availability, or between *ABO* blood type variants and several different bacterial families depending on the cohort (3). In turn, studies in mice comparing gut microbiome composition between wildtype mice and loss-of-function mutants revealed contributions of several innate immune related genes to determining the composition of the gut microbiome (4–7). However, the role of host genes in determining microbiome composition is sometimes not immediately discernable in mouse mutants, requiring several generations to become evident, which in some cases was interpreted to be the result of drift rather than the mutation itself, although in other cases such “drift” was subsequently shown to be indeed due to accumulating effects of candidate gene disruptions (8–10).

Invertebrate model organisms such as *Drosophila melanogaster* and *Caenorhabditis elegans* offer alternative models with greater genetic tractability, and similar to vertebrates, have demonstrated the importance of host immunity for controlling gut microbiome composition (11–14). Work in drosophila further revealed differential activation of immune mechanisms by pathogens or by non-pathogenic gut commensals, highlighting the ability of the innate immune system (which drosophila, as all other invertebrates, solely rely on) to provide variable responses to maintain homeostasis and prevent collateral damage (11, 15). Work with age-synchronized populations of *C. elegans* in turn demonstrated how an age-dependent decline in a pathway of immune control was associated with age-dependent dysbiosis, and the importance of a diverse gut community for preventing the detrimental consequences of this dysbiosis (16).

‘Common garden’ experiments, in which different *C. elegans* strains and related species were raised in identical compost microcosms, identified a significant contribution of host genetics to determining microbiome composition (17). Subsequent studies identified conserved regulatory pathways, including insulin/insulin-like (IIS) signaling (18, 19) and the DBL-1/BMP pathway (12), as contributing to shaping of the gut microbiome. DBL-1 signaling further came to the forefront as a mechanism that controls a specific subset of gut bacteria, which has the potential to cause detrimental effects when control was impaired (12). The DBL-1 ligand, a BMP-1 homolog, is primarily expressed in neurons (20), and upon secretion activates a broadly expressed heterodimer receptor, and downstream to it drives nuclear localization of transcriptional regulators SMA-2, –3 and – 4, to activate gene expression (21). While DBL-1 signaling contributes both to larval development and to immunity, its effects on the gut microbiome were linked specifically to its immune contributions (12). Disruption of genes for any of the DBL-1 pathway’s components led to an expansion specifically of gut bacteria of the *Enterobacteriaceae* family, particularly of the genus *Enterobacter*. However, experiments attempting to rescue DBL-1 control in *sma-3* mutants, through tissue specific *sma-3* expression, revealed that control over gut *Enterobacter* could not be achieved through intestinal *sma-3* expression, and that, instead, expression from the epidermis or pharynx could restore control, suggesting that DBL-1 and SMA-3 signaling affected the gut microbiome cell non-autonomously, likely dependent on downstream activation of intestinal mediators.

Contributions of central regulatory pathways to shaping of the gut microbiome are large and thus easier to detect. Identifying smaller contributions of each individual downstream effector is more of a challenge. To understand how DBL-1 signaling affected the gut microbiome, we carried out RNA-seq analysis and subsequent functional characterization of candidate mediators, which led to identification of several DBL-1-regulated intestinal effectors with potential additive contributions to control of *Enterobacteriaceae* gut colonization. This expands our understanding of the contributions of DBL-1 signaling to describe a gene network operating downstream to it, which contributes to shaping of the gut microbiome.

## RESULTS

### Targets of DBL-1/BMP signaling include microbiome-modulated immune genes

To identify genes regulated by DBL-1/BMP signaling in the context of interactions with a complex microbial community, we performed RNA-seq analysis comparing gene expression in adult wildtype worms, *dbl-1* and *sma-3* mutants, and *dbl-1* over-expressing transgenics, raised either on non-colonizing *E. coli* or on the CeMbio community (22). Sleuth analysis identified 2291 genes differentially expressed in DBL-1/BMP-perturbed strains (*q* < 0.005), divided between four clusters with distinct expression patterns (Figure 1A, Supplementary Data 1). Cluster 1 included 742 genes that were upregulated to a varying extent on CeMbio, less so in either one of the two mutant strains, and much more in *dbl-1* over-expressing worms; Cluster 2 included 503 genes, which while also dependent for their basal expression on DBL-1 signaling (lower in mutants, higher in over-expressing animals), were repressed on CeMbio. Analysis of enriched annotations revealed enrichment for immune and stress response genes in both clusters, including C-type lectins and genes involved in detoxification, supporting the role of the DBL-1 pathway in immune regulation. However, differences in gene composition between the two clusters were also apparent, with the CeMbio-upregulated genes of Cluster 1 showing a prominent enrichment for C-type lectins, while the CeMbio-downregulated genes of Cluster 2, showed more significant enrichment for detoxification genes, suggesting that DBL-1 signaling contributed differentially to the expression of different subsets of immune and stress genes. Cluster 1 further featured a significant enrichment for genes previously identified to be induced in response to two different complex communities (6 of 30 genes, *p* < 0.001, hypergeometric test, Supplementary table 1) (12). Cluster 4 was of additional interest, including 844 genes that were negatively regulated by DBL-1 signaling. Among them, enrichment was found for genes involved in house-keeping functions, such as mRNA processing (e.g. *prp-x/xx*, *rnp-x/xx*) and protein synthesis (e.g. *rps-x/xx* and *rpl-x*), suggesting the involvement of DBL-1 signaling in negative regulation of growth and maintenance functions in adults, in contrast to its better known positive contributions to cell growth in larvae (23).

**Figure 1.**
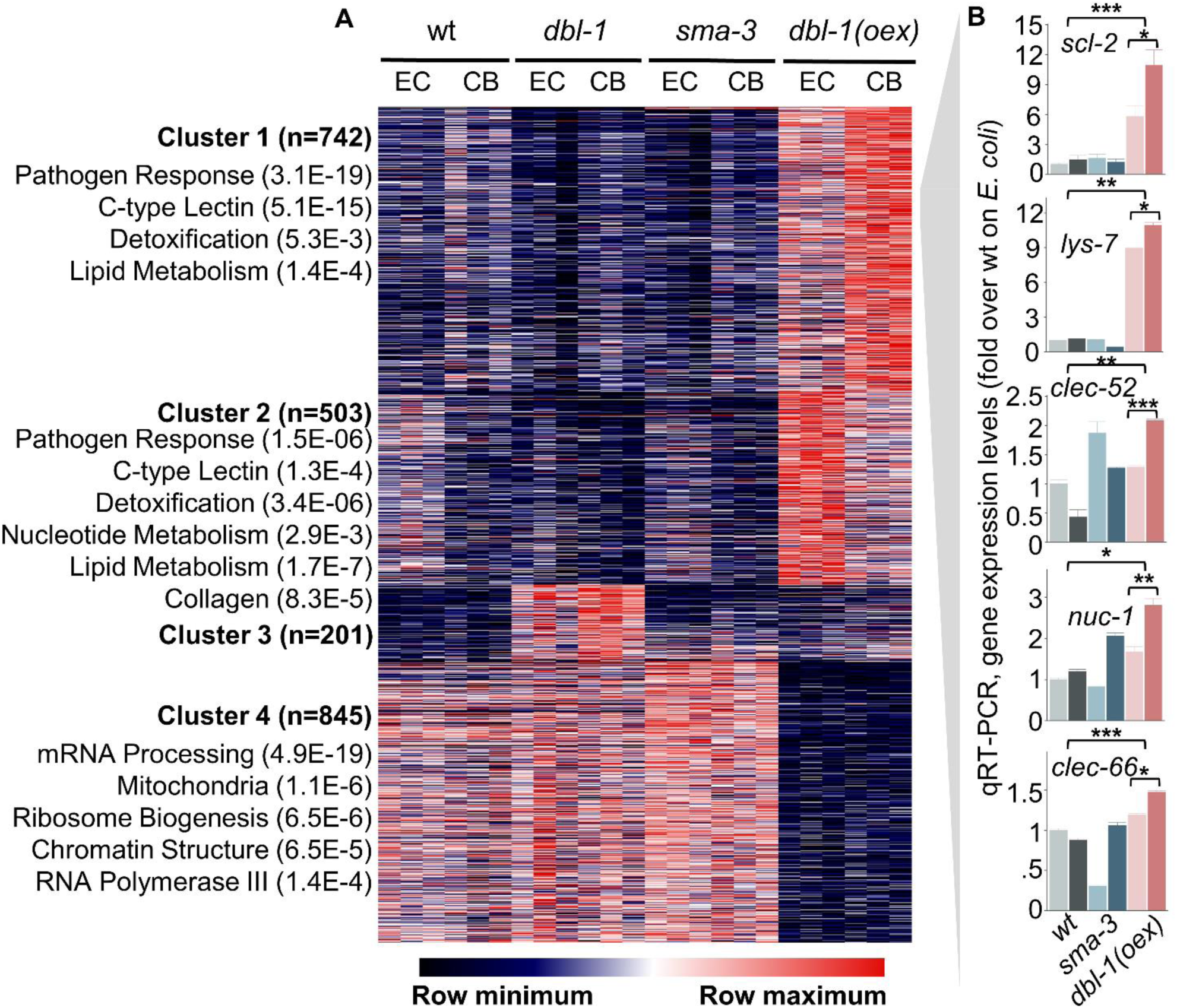
DBL-1/BMP-dependent gene expression. (**A**) Expression profiles of Sleuth-identified DBL-1 pathway targets (p < 0.005, BH-corrected, see methods) in wildtype and designated mutant and transgenic strains raised on *E. coli* (EC) or on CeMbio (CB). Genes are *k*-means-clustered (with number of genes for each cluster) and colored following median-centering for each gene to highlight patterns. Enriched gene annotations were identified using WormCat, with Bonferroni-corrected p-values **(B)** qRT-PCR verification of expression patterns for putative DBL-1 pathway targets of Cluster 1 in the designated strains; light and dark colors represent expression in worms raised on *E. coli* or CeMbio, respectively. Shown for each graph are averages of two independent experiments (N = 2) ± SDs. * *p* < 0.05, ** *p* < 0.01, *** *p* < 0.001, pairwise t-test.

Focusing on genes of Cluster 1 – positively regulated by DBl-1 signaling and upregulated in response to CeMbio – we selected five, *scl-2*, *lys-7*, *clec-52 nuc-1,* and *clec-66* (of which the first four were previously identified to be upregulated by complex communities (12)) for additional analyses using qRT-PCR. Overall, qRT-PCR measurements supported the identification of these genes as regulated by DBL-1, most clearly seen in the *dbl-1* over-expressing strain (Fig. 1B). However, only *lys-7* and *clec-66* showed some indication of reduced expression in *sma-3* mutants, suggesting that identified DBL-1 targets receive additional regulatory inputs that could keep their expression at normal levels in *sma-3* mutants. Indeed, *lys-7*, *clec-52* and *nuc-1* were previously reported to be regulated also by the longevity and immune-associated transcription factor DAF-16 (24), and *lys-7* and *clec-52* were reported also as targets of the stress activated p38 MAPK pathway (25, 26). Thus, these genes appear to be regulated redundantly, with DBL-1 signaling being one of several regulatory inputs.

### Involvement of DBL-1 targets in determining gut microbiome composition

The five verified DBL-1 targets are known to be expressed in the intestine (wormbase.org) and all contain signal peptides targeting for their secretion (27), suggesting that they could interact directly with gut bacteria. *scl-2* encodes a yet uncharacterized protein homologous to mammalian cysteine-rich secreted proteins and peptidase inhibitors, which are best characterized for their ability to coat sperm cells to facilitate fertilization (28); *lys-7* encodes a lysozyme with documented roles in anti-bacterial defense (29); *clec-52* (ortholog of human Reg3a) and *clec-66* encode C-type lectins, thought to bind bacterial surface saccharides (30, 31); and lastly, *nuc-1* encodes a nuclease that degrades apoptotic DNA (32), which was additionally reported to digest bacterial DNA in the intestine (33).

Previously, we identified the role of DBL-1 signaling in regulating the colonization of *Enterobacter hormaechei* strain CEent1. To determine if the presently-identified five DBL-1 targets may serve as downstream mediators of this interaction, we tested the level of colonization of a fluorescently tagged derivative of CEent1 in mutant strains for the five DBL-1 targets (12, 16). Among worms raised on monocultures of CEent1-dsRed, significantly increased colonization was observed in four of the examined mutants compared to wildtype worms (excluding *clec-52*), but for the most part the extent of increase was lower than in *sma-3* mutants, supporting the candidate genes’ involvement in mediating the contributions of DBL-1 signaling to control of gut bacterial colonization (Fig. 2). Interestingly, *nuc-1* mutants showed exceptionally increased colonization, greater than that seen in *sma-3*. To test how the disruption of the candidate genes may affect a more complex gut community rather than a single colonizer, we raised wildtype and mutant worms on the CeMbio community of twelve strains and analyzed their gut microbiome composition using V4 16S sequencing. This analysis identified significant differences between wildtype animals and most mutants, excluding *clec-66* (Fig. 3A). Gut microbiomes assembled from CeMbio tend to be dominated by two strains – *Ochrobactrum vermis* (MYb71) and *Stenotrophomonas indicatrix* (JUb19) – contributing 70-80% of total bacterial abundance (22), and this dominance was maintained in the examined mutants (Supplementary Data 2). However, relative abundance of *Enterobacteriaceae* strains *E. hormaechei* (CEent1) and *Lelliottia amnigena* (JUb66), which cannot be distinguished based on V4 16S sequencing, increased reproducibly in four out of five mutants (excluding *lys-7*), extending the previously described role of DBL-1 signaling in control of members of the *Enterobacteriaceae* family to its putative downstream mediators (Fig. 3B and C) (12). Additional experiments were performed to complement the sequencing analysis of gut microbiome composition, using CFU counts of gut bacteria isolated from wildtype and mutant worms raised on CeMbio. Samples were split between rich media and *Enterobacteriaceae*-selective VRBG media plates, to assess total bacterial load, or *Enterobacteriaceae* load, respectively. While total bacterial load did not change significantly in most mutants compared to wildtype animals (Fig. 4A), the proportion of *Enterobacteriaceae* increased significantly in most examined mutants, excluding *clec-66* (Fig. 4B and C). Together, the results from these different experimental techniques support the involvement of *scl-2* and *nuc-1* in controlling *Enterobacteriaceae* gut abundance in both monocultures and in the context of the CeMbio community, with *lys-7*, *clec-52* and *clec-66* showing smaller and less reproducible contributions.

**Figure 2.**
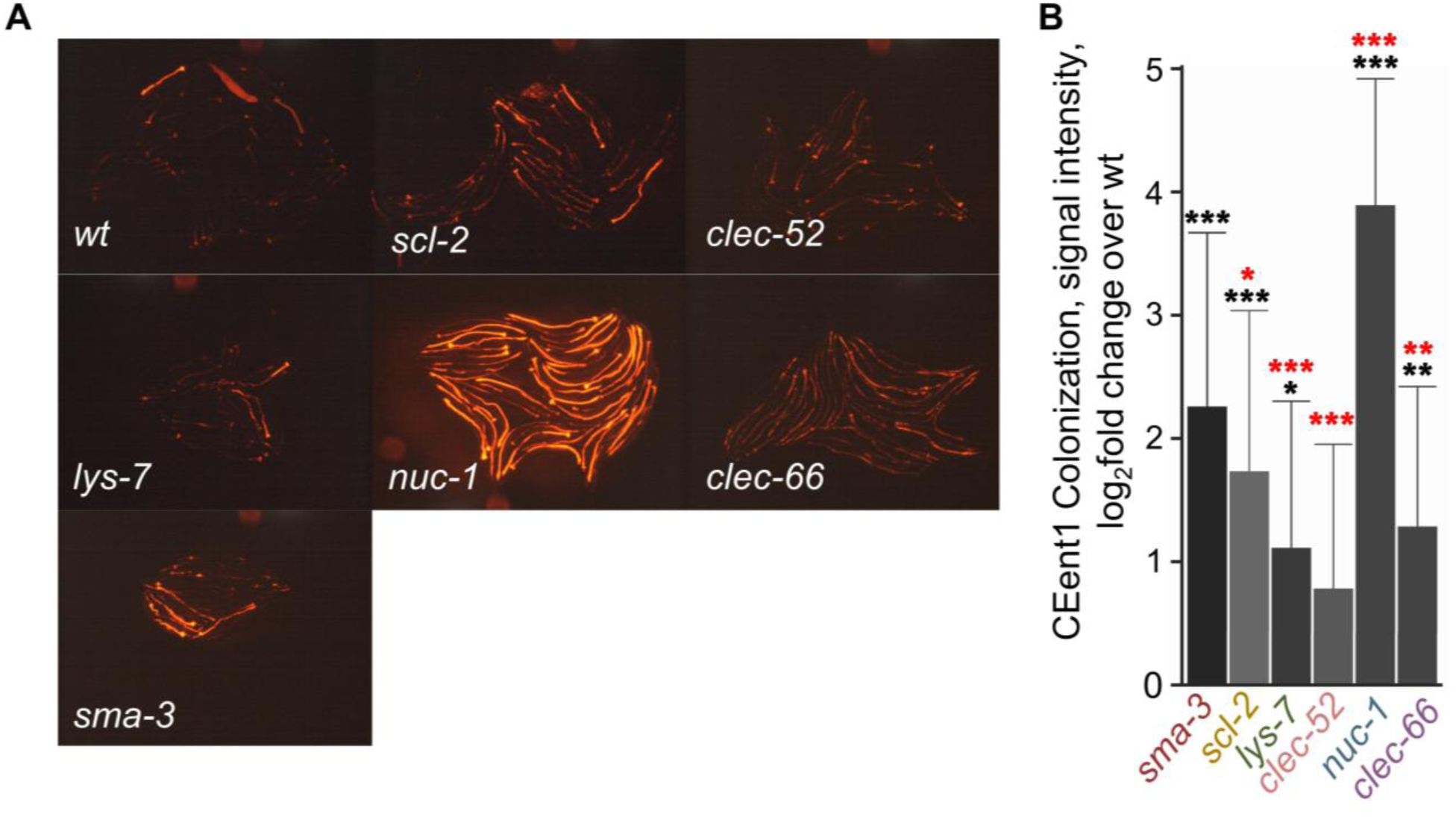
Disruption of intestinal DBL-1 target genes increases *Enterobacter* colonization. (**A**) Representative images of DBL-1/BMP effector mutant strains grown on CEent-1-dsRed bacteria, compared to wildtype (wt), recorded 1 d after L4. Scale bar, 200 μm. **(B)** Quantification of signal intensity in worms as in A. Bars mark average single worm fluorescence + SDs; 20-46 worms per experiment (N=4 independent experiments for *scl-2* and *clec-52*; N=3, for *lys-7* and *clec-66*; and N=2 2, for *nuc-1*); * *p* < 0.05, *** *p* < 0.001, t-test compared to wt; red for comparison to *sma-3*.

**Figure 3.**
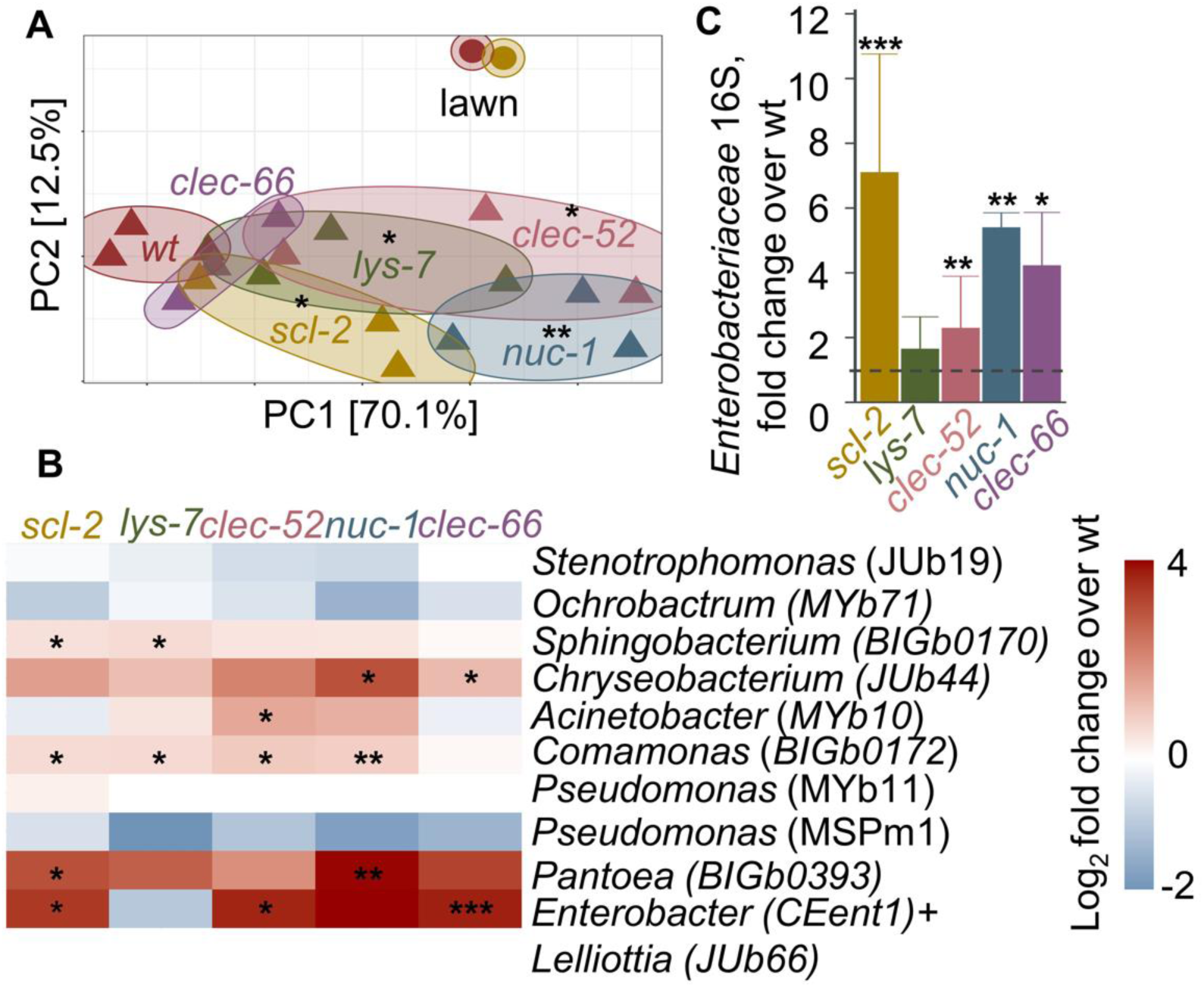
Disruption of intestinal DBL-1 targets alters gut microbiome composition. (**A**) PCoA based on weighted UniFrac distances highlighting differences in microbiome composition (analyzed by 16S NGS) between wildtype and mutant worms in one experiment, analyzed for each strain in three independent populations; * *p* < 0.05, ** *p* < 0.01, UniFrac regression-based kernel association test with small-sample size correction. **(B)** Data from A, highlighting relative abundances of CeMbio members in tested mutants, shown as fold over wildtype; * *p* < 0.05, ** *p* < 0.01, *** *p* < 0.001, t-test, compared to wildtype. (**C**) *Enterobacteriaceae* relative abundance in designated mutants, including results from several independent experiments as the one presented in A (N = 4 for *scl-2* and *clec-52* mutants, N = 2 for *nuc-1* and *clec-66,* and N = 1 for *lys-*7), each performed with 3-5 worm populations. Values are shown as fold over wildtype, * *p* < 0.05, ** *p* < 0.01, *** *p* < 0.001, t-test compared to wildtype.

**Figure 4.**
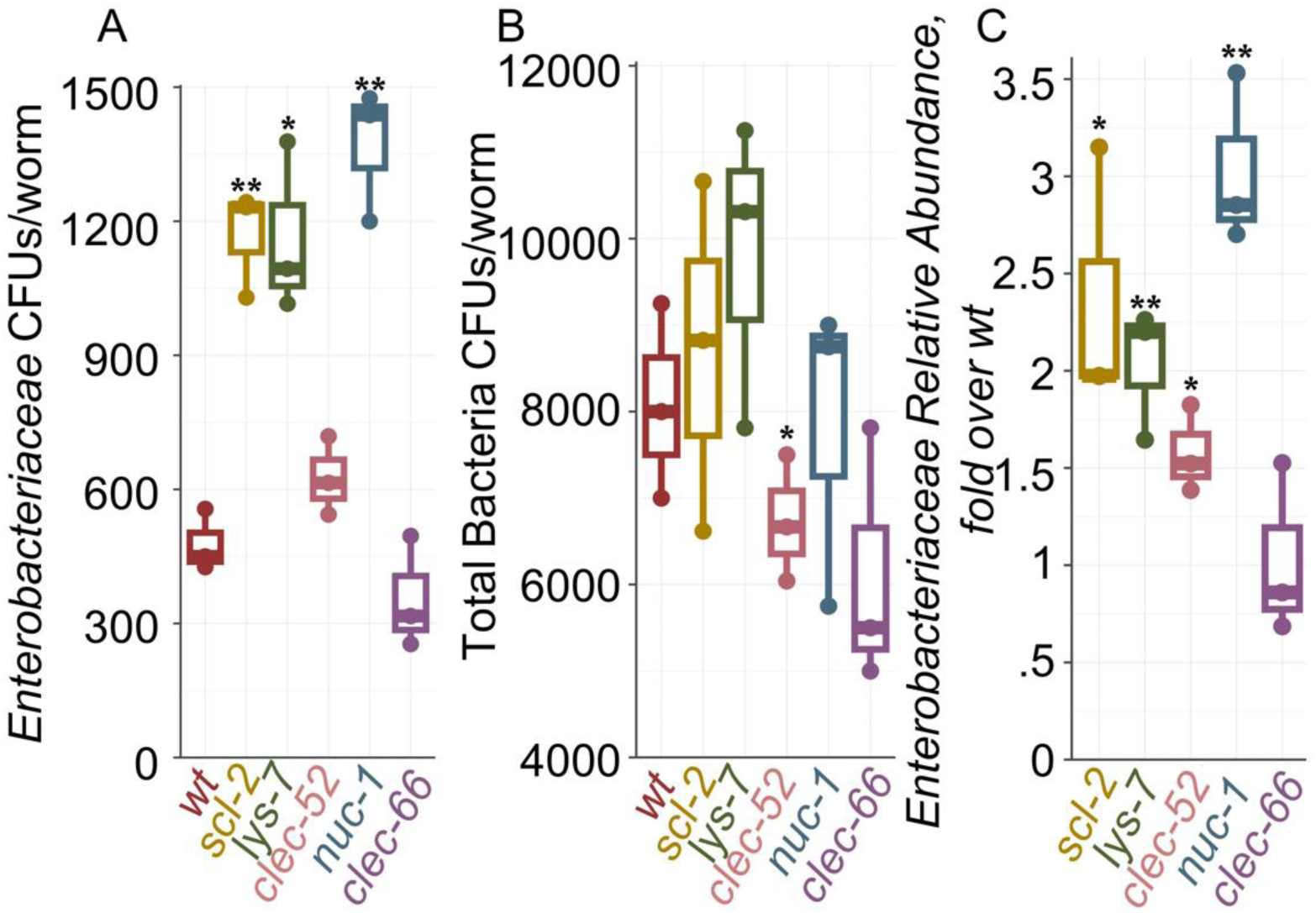
Disruption of intestinal DBL-1 targets increases gut *Enterobacteriaceae* load. (**A**) *Enterobacteriaceae* gut load, represented by CFU counts on selective VRBG media. (B) Total bacterial load, counted on LB plates. (C) *Enterobacteriaceae* proportion of total bacterial load relative to wildtype calculated from data in A and B. * *p* < 0.05, ** *p* < 0.01, *** *p* < 0.001, t-test compared to wildtype.

## DISCUSSION

Previous identification of DBL-1/BMP immune signaling as a factor determining gut microbiome composition, specifically controlling abundance of *Enterobacteriaceae*, raised the question of what mediated its effects on gut bacteria. DBL-1-dependent control was deemed to involve several regulatory levels, as its effects depended on activation of transcriptional regulators in extra-intestinal tissues (12, 34). The results described here begin to fill-in this gap by identifying several intestinal mediators, likely secreted, which could directly interact with gut bacteria to control their abundance. The five examined mediator genes, *scl-2*, *lys-7*, *clec-52*, *clec-66,* and *nuc-1* are positively regulated by DBL-1 signaling, but very likely not directly, as they were upregulated by *dbl-1* overexpression, but not downregulated by *dbl-1* disruption, suggesting additional regulatory inputs. For some, i.e. *lys-7*, *clec-52* and *nuc-1,* such inputs – by DAF-16 and p38 MAPK signaling, were previously described, and may also be responsible for relaying extra-intestinal DBL-1 signaling to activate intestinal mediators (24–26). Although to a varying extent and reproducibility, disruption of each of the five putative mediators led to increases in the relative abundance of species of the *Enterobacteriaceae* family and in the actual number of such cells in the gut, supporting their importance for gut bacterial control.

The lysozyme gene and the C-type lectin genes are known to be associated with responses to pathogenic bacteria (35–38). Our results extend their function to controlling non-pathogenic commensal bacteria. At least for one of these genes, *clec-52,* this involvement may be conserved, as enteric delivery of the human *clec-52* homolog Reg3A in mice was shown to alter gut microbiome composition and to reduce colitis (39). In contrast, *scl-2* and *nuc-1* are not typically associated with immune responses, but are consistently found to be upregulated in worms exposed to complex microbial communities (12), supporting their involvement in host-microbiome interactions.

The experiments presented in Fig. 2, following worm gut colonization with CEent1-dsRed, demonstrated that with the exception of *nuc-1*, the effect of disrupting each of the putative mediator genes was smaller than that seen in worms disrupted for the upstream regulator gene, *sma-3*. This supports the hypothesis that control of *Enterobacter* colonization by DBL-1 signaling relies on a cocktail of downstream effectors, each with a small contribution and together accumulating to the full effect seen in *sma-3* mutants. Other regulatory pathways may induce the expression of other antimicrobial cocktails, partially overlapping in their composition to those regulated by DBL-1 and affect non-*Enterobacteriaceae* gut bacteria. For example, insulin signaling (IIS), mediated by DAF-16 (which contributes also to the expression of some of the DBL-1 targets identified here), was shown to control abundance of bacteria of the genus *Ochrobactrum*, also common inhabitants of the worm gut (18). Through partially overlapping antimicrobial cocktails, a few regulatory pathways could differentially control gut microbes and shape microbiome composition. Several studies, primarily in drosophila, demonstrate the contributions of different immune regulators to the abundance of different gut constituents (11). A recent study, also in drosophila, nicely demonstrated differential control, describing specific effects of Diptericin A and B, two antimicrobial peptides regulated by the Imd immune pathway, on two distinct gut commensals (40). This observation further suggests that diversification of antimicrobial proteins may be driven not only by the need to fight pathogens but also by the need to control gut microbiome composition.

Unlike most of the examined mediators, *nuc-1* disruption led to a dramatic increase in *Enterobacteriaceae* abundance, larger than seen in its *sma-3* regulator. *nuc-1*, encoding a DNase II homolog, is thought to be important for degradation of DNA in cells undergoing apoptosis (32). It has been also reported to be involved in degradation of bacterial DNA in the intestinal lumen (33), but this suggestion could not explain the effects of its disruption of the increase of intact gut bacteria, as observed in analysis of colony forming units. Another study suggested that *nuc-1* disruption in the germline upregulated antimicrobial peptides (41), but this again could not explain the increase in gut bacteria that we observed. Thus, while *nuc-1* appears to play an important role in controlling gut bacteria, at this point, the mechanism remains unknown.

While the mechanisms underlying the effects of the identified intestinal mediators on the gut microbiome remain to be investigated, our results describe a new layer in worm control over its gut bacteria and expand our understanding of the role of DBL-1 signaling in such control to describe an underlying gene network that mediates its effects on the worm gut microbiome.

## METHODS

**Worm strains used** in this study included the N2 wildtype strain, *dbl-1(nk3), sma-3(e491)*, and the *dbl-1* overexpressing strain BW1940[*dbl-1p::dbl-1;sur-5::gfp*] (20), *lys-7(ok1384)*, *nuc-1(e1392)*, and *clec-66(ok2230)*, all obtained from the *Caenorhabditis* Genome Center (CGC), and *clec-52(tm8126)* and *scl-2(tm2428),* obtained from the National Bioresource Project (42). Worms were raised on standard nematode growth medium (NGM) or on peptone-free medium (43), with bacteria as food or as colonizers.

**Bacterial strains and communities** included the non-colonizing *E. coli* strain OP50, used as food and as control, CeMbio (22), a defined community of *C. elegans* gut commensals consisting of twelve characterized strains selected to represent the core *C. elegans* gut microbiome, and CEent1-dsRed, a fluorescently-tagged derivative of the *Enterobacter hormachei* strain CEent1, a member of the CeMbio community (16). CeMbio strains were raised as previously described (22), normalized based on optical density, mixed in equal proportions and seeded on appropriate plates.

**RNA-seq**. Germ-free L1 larvae obtained from gravid worms by bleaching (three independent populations per worm strain) were raised at 25°C on NGM plates seeded with CeMbio as described above. Gravid worms were rinsed off plates with M9 including 0.025% Triton, washed 5 times to get rid of offspring and bacteria, mixed with TRIzol Reagent (Invitrogen; Waltham, USA), snap-frozen in liquid nitrogen, taken through 5-7 thaw-freeze cycles to break them open, and kept at –80°C until use. RNA isolation was performed using the NucleoSpin RNA purification kit, manual protocol 5.2 (Macherey-Nagel; Düren, Germany).

Sequencing libraries were prepared from total RNA using the TruSeq RNA Library Kit v2 (Illumina; San Diego, USA), with indexed adaptors for multiplex sequencing, assessed for quality on an Agilent Bio-analyzer (Agilent; Santa Clara, USA) and submitted for 100 bp paired-end sequencing on a NovaSeq 6000 at the QB3 Genomic Sequencing Laboratory (UC Berkeley, Berkeley, CA; RRID:SCR_022170). Raw reads obtained were pre-processed with *fastp* (44) and pseudo-aligned to the WormBase transcriptome version WS235 using *kallisto* (45). Transcript counts were then normalized with *Sleuth* (46) and analyzed to identify genes differentially expressed between worm strains and bacterial treatment using the likelihood ratio test. Heatmaps following *k*-means clustering (*k* = 4) were generated with *Morpheus* (https://software.broadinstitute.org/morpheus) and gene set enrichment analyses were performed using WormCat (47).

**Quantitative (q)RT-PCR** measurements were performed on RNA extracted as described above from worms raised at 20°C as described above. mRNA was reverse transcribed with the iScript Reverse Transcription Supermix (BioRad, Hercules, USA), and cDNA was used for amplification using the SsoAdvanced Universal SYBR Green Supermix (BioRad, Hercules, USA) on an Applied Biosystems StepOnePlus cycler (Waltham, USA). Ct values obtained in amplification of specific mRNAs were normalized to those obtained by amplification of three conserved *C. elegans* actin genes with the pan-actin primer pair (35).

Primers used included:

*scl-2*: F: 5’-GATTTCGCCCACGCCATTTG-3’; R: 5’-ACTCAGAAATCGCCGGGAAC –3’

*lys-7*: F 5’-TTGCAGTACTCTGCCATTCG-3’; R: 5’-GCACAATAACCCGCTTGTTT –3’

*clec-52*: F: 5’-AGCCAAATCTCCTCCATCAGC-3’; R: 5’-GATCAACCGCCTGTATGCAAC –3’

*nuc-1*: F: 5’-CCTGGAAGATGGTCTTGTCA-3’; R: 5’-GGGAACTTTGACTCCTTCTGC –3’

*clec-66*: F: 5’-GCAGAAGGCGGTTTTGGC-3’; R: 5’-GCGGCGAATTTAGTCATGGC –3’

PanActin: F: 5’-TCGGTATGGGACAGAAGGAC-3’; R: 5’-CATCCCATGTGGTGACGATA –3’

**DNA extraction for gut microbiome analysis.** Gravid worms raised at 20°C on NGM plates with CeMbio (three independent populations per worm strain) were washed off plates, washed 5 times with M9 + 0.025% Triton, paralyzed with levamisole to seal the intestine, surface sterilized with bleach as described elsewhere (22, 48), and kept at 4 °C until use. DNA was extracted using the Qiagen DNeasy PowerSoil Pro Kit, with modifications as described elsewhere (48).

**16S rRNA gene sequencing** of the amplicon libraries of the V4 variable region generated with primers 515F and 806R containing Illumina overhang adapter sequences according to the manufacturer instructions, with slight cycling modifications described elsewhere. Dual indices and Illumina sequencing adapters were added using the Nextera XT Index Kit. Sequencing was performed on an Illumina MiniSeq.

Demultiplexed forward and reverse sequences were filtered for quality, resulting in roughly 11,000 reads per sample, and assigned amplicon sequence variants (ASVs) with DADA2 (49) and *DECIPHER* (50). Taxonomy assignments for ASVs were obtained based on a custom database with 16S sequences of the twelve CeMbio strains, and counts were normalized for the different 16S gene copy number of the different strains. Microbiome analyses were performed in R using *phyloseq* (51), *phangorn* (52), and *vegan* (53), to calculate UniFrac distances for Principle Component Analysis; and MiRKAT (54), for statistical evaluation of differences between microbiomes.

**Colony forming unit (CFU) Counts** of gut commensals were evaluated in worms raised, harvested and surface-sterilized as described above. Gut bacteria were released from worms by vortexing together with zirconium beads, until degradation could be confirmed using a light microscope. Serially diluted worm lysates were plated on either *Enterobacteriaceae*-selective media (Violet Red Bile Glucose, VRBG; Difco Becton Dickinson) or on rich LB media and incubated at 28°C for 24 h before counting colonies.

**Fluorescence Imaging** was employed to follow worm colonization by *E. hormachei* CEent1, using the CEent1-dsRed derivative. Worms were raised from the L1 stage on a lawn of CEent-1-dsRed at 20°C. Following three days, gravid worms were washed off plates, washed three times with M9 and imaged. Fluorescent images were captured using a Leica MZ16F equipped with a QImaging MicroPublisher 5.0 camera and fluorescent signal of colonizing bacteria was quantified on the Fiji plugin of ImageJ v2.10/1.53c as previously described (55), producing background-subtracted average intensity for each worm.

## Acknowledgements

Research toward this manuscript was made possible thanks to NIH grant 1R01OD024780-01A1. K.T. was supported by summer fellowships funded by the Rose Hills Foundation and the Office of Undergraduate Research & Scholarship at UC Berkeley. Some strains were provided by the CGC, which is funded by NIH Office of Research Infrastructure Programs (P40 OD010440).

## Author Contributions

BP and MS conceived the project; KT, BP, SK, assisted by RB and SH, performed all experiments, and analyzed their results. KT and MS wrote the manuscript, with edits from BP.

## Data availability

Raw RNA-seq data and *kallisto* output files have been deposited in GEO with accession number GSE186653; the associated computational pipeline is available online at https://github.com/rahulnccs/TGF-beta_RNAseq_Analysis. 16S sequencing data is available in the NCBI Sequence Read Archive (Bioproject ID PRJNA1031602), with the computational pipeline available at https://github.com/kennytrang/DBL-1_Mediators.

## References

1. Gilbert JA, Lynch SV. 2019. Community ecology as a framework for human microbiome research. Nat Med 25:884–889.

2. Ryu EP, Davenport ER. 2022. Host Genetic Determinants of the Microbiome Across Animals: From Caenorhabditis elegans to Cattle. Annu Rev Anim Biosci 10:203–226.

3. Sanna S, Kurilshikov A, Van Der Graaf A, Fu J, Zhernakova A. 2022. Challenges and future directions for studying effects of host genetics on the gut microbiome. Nat Genet 54:100–106.

4. Lamas B, Richard ML, Leducq V, Pham H-P, Michel M-L, Da Costa G, Bridonneau C, Jegou S, Hoffmann TW, Natividad JM, Brot L, Taleb S, Couturier-Maillard A, Nion-Larmurier I, Merabtene F, Seksik P, Bourrier A, Cosnes J, Ryffel B, Beaugerie L, Launay J-M, Langella P, Xavier RJ, Sokol H. 2016. CARD9 impacts colitis by altering gut microbiota metabolism of tryptophan into aryl hydrocarbon receptor ligands. Nat Med 22:598–605.

5. Salzman NH, Hung K, Haribhai D, Chu H, Karlsson-Sjöberg J, Amir E, Teggatz P, Barman M, Hayward M, Eastwood D, Stoel M, Zhou Y, Sodergren E, Weinstock GM, Bevins CL, Williams CB, Bos NA. 2010. Enteric defensins are essential regulators of intestinal microbial ecology. Nat Immunol 11:76–82.

6. Vijay-Kumar M, Aitken JD, Carvalho FA, Cullender TC, Mwangi S, Srinivasan S, Sitaraman SV, Knight R, Ley RE, Gewirtz AT. 2010. Metabolic Syndrome and Altered Gut Microbiota in Mice Lacking Toll-Like Receptor 5. Science 328:228–231.

7. Petnicki-Ocwieja T, Hrncir T, Liu Y-J, Biswas A, Hudcovic T, Tlaskalova-Hogenova H, Kobayashi KS. 2009. Nod2 is required for the regulation of commensal microbiota in the intestine. Proc Natl Acad Sci 106:15813–15818.

8. Viennois E, Pujada A, Sung J, Yang C, Gewirtz AT, Chassaing B, Merlin D. 2020. Impact of PepT1 deletion on microbiota composition and colitis requires multiple generations. Npj Biofilms Microbiomes 6:27.

9. Lemire P, Robertson SJ, Maughan H, Tattoli I, Streutker CJ, Platnich JM, Muruve DA, Philpott DJ, Girardin SE. 2017. The NLR Protein NLRP6 Does Not Impact Gut Microbiota Composition. Cell Rep 21:3653–3661.

10. Mamantopoulos M, Ronchi F, Van Hauwermeiren F, Vieira-Silva S, Yilmaz B, Martens L, Saeys Y, Drexler SK, Yazdi AS, Raes J, Lamkanfi M, McCoy KD, Wullaert A. 2017. Nlrp6– and ASC-Dependent Inflammasomes Do Not Shape the Commensal Gut Microbiota Composition. Immunity 47:339–348.e4.

11. Lesperance DN, Broderick NA. 2020. Microbiomes as modulators of Drosophila melanogaster homeostasis and disease. Curr Opin Insect Sci 39:84–90.

12. Berg M, Monnin D, Cho J, Nelson L, Crits-Christoph A, Shapira M. 2019. TGFβ/BMP immune signaling affects abundance and function of C. elegans gut commensals. Nat Commun 10:604.

13. Zhang F, Berg M, Dierking K, Félix M-A, Shapira M, Samuel BS, Schulenburg H. 2017. Caenorhabditis elegans as a Model for Microbiome Research. Front Microbiol 8.

14. Shapira M. 2017. Host–microbiota interactions in Caenorhabditis elegans and their significance. Curr Opin Microbiol 38:142–147.

15. Bosch TCG, Zasloff M. 2021. Antimicrobial Peptides—or How Our Ancestors Learned to Control the Microbiome. mBio 12:e01847–21.

16. Choi R, Bodkhe R, Pees B, Kim D, Berg M, Monnin D, Cho J, Narayan V, Deller E, Shapira M. 2023. An Enterobacteriaceae bloom in aging animals is restrained by the gut microbiome. bioRxiv 10.1101/2023.06.13.544815.

17. Berg M, Zhou XY, Shapira M. 2016. Host-Specific Functional Significance of Caenorhabditis Gut Commensals. Front Microbiol 7.

18. Zhang F, Weckhorst JL, Assié A, Hosea C, Ayoub CA, Khodakova AS, Cabrera ML, Vidal Vilchis D, Félix M-A, Samuel BS. 2021. Natural genetic variation drives microbiome selection in the Caenorhabditis elegans gut. Curr Biol 31:2603–2618.e9.

19. Taylor M, Vega NM. 2021. Host Immunity Alters Community Ecology and Stability of the Microbiome in a Caenorhabditis elegans Model. mSystems 6:e00608–20.

20. Suzuki Y, Yandell MD, Roy PJ, Krishna S, Savage-Dunn C, Ross RM, Padgett RW, Wood WB. 1999. A BMP homolog acts as a dose-dependent regulator of body size and male tail patterning in Caenorhabditis elegans. Development 126:241–250.

21. Savage-Dunn C. 2001. Targets of TGFβ-related signaling in Caenorhabditis elegans. Cytokine Growth Factor Rev 12:305–312.

22. Dirksen P, Assié A, Zimmermann J, Zhang F, Tietje A-M, Marsh SA, Félix M-A, Shapira M, Kaleta C, Schulenburg H, Samuel BS. 2020. CeMbio – The Caenorhabditis elegans Microbiome Resource. G3 Genes Genomes Genetics 10:3025–3039.

23. Savage C, Das P, Finelli AL, Townsend SR, Sun CY, Baird SE, Padgett RW. 1996. Caenorhabditis elegans genes sma-2, sma-3, and sma-4 define a conserved family of transforming growth factor beta pathway components. Proc Natl Acad Sci 93:790– 794.

24. Murphy CT, McCarroll SA, Bargmann CI, Fraser A, Kamath RS, Ahringer J, Li H, Kenyon C. 2003. Genes that act downstream of DAF-16 to influence the lifespan of Caenorhabditis elegans. 6946. Nature 424:277–283.

25. Fletcher M, Tillman EJ, Butty VL, Levine SS, Kim DH. 2019. Global transcriptional regulation of innate immunity by ATF-7 in C. elegans. PLOS Genet 15:e1007830.

26. Troemel ER, Chu SW, Reinke V, Lee SS, Ausubel FM, Kim DH. 2006. p38 MAPK Regulates Expression of Immune Response Genes and Contributes to Longevity in C. elegans. PLoS Genet 2:e183.

27. Suh J, Hutter H. 2012. A survey of putative secreted and transmembrane proteins encoded in the C. elegans genome. BMC Genomics 13:333.

28. Gibbs GM, Roelants K, O’Bryan MK. 2008. The CAP Superfamily: Cysteine-Rich Secretory Proteins, Antigen 5, and Pathogenesis-Related 1 Proteins—Roles in Reproduction, Cancer, and Immune Defense. Endocr Rev 29:865–897.

29. Boehnisch C, Wong D, Habig M, Isermann K, Michiels NK, Roeder T, May RC, Schulenburg H. 2011. Protist-Type Lysozymes of the Nematode Caenorhabditis elegans Contribute to Resistance against Pathogenic Bacillus thuringiensis. PLoS ONE 6:e24619.

30. Pees B, Yang W, Kloock A, Petersen C, Peters L, Fan L, Friedrichsen M, Butze S, Zárate-Potes A, Schulenburg H, Dierking K. 2021. Effector and regulator: Diverse functions of C. elegans C-type lectin-like domain proteins. PLOS Pathog 17:e1009454.

31. Pees B, Yang W, Zárate-Potes A, Schulenburg H, Dierking K. 2016. High Innate Immune Specificity through Diversified C-Type Lectin-Like Domain Proteins in Invertebrates. J Innate Immun 8:129–142.

32. Wu Y-C, Stanfield GM, Horvitz HR. 2000. NUC-1, a Caenorhabditis elegans DNase II homolog, functions in an intermediate step of DNA degradation during apoptosis. Genes Dev 14:536–548.

33. Lai H-J, Lo SJ, Kage-Nakadai E, Mitani S, Xue D. 2009. The Roles and Acting Mechanism of Caenorhabditis elegans DNase II Genes in Apoptotic DNA Degradation and Development. PLOS ONE 4:e7348.

34. Choi R, Kim D, Li S, Massot M, Narayan V, Slowinski S, Schulenburg H, Shapira M. 2020. Extra-intestinal regulation of the gut microbiome: The case of C. elegans TGFβ/SMA signaling, p. 119–134. In Cellular Dialogues in the Holobiont. CRC Press.

35. Shapira M, Hamlin BJ, Rong J, Chen K, Ronen M, Tan M-W. 2006. A conserved role for a GATA transcription factor in regulating epithelial innate immune responses. Proc Natl Acad Sci 103:14086–14091.

36. Mallo GV, Kurz CL, Couillault C, Pujol N, Granjeaud S, Kohara Y, Ewbank JJ. 2002. Inducible Antibacterial Defense System in C. elegans. Curr Biol 12:1209–1214.

37. Evans EA, Kawli T, Tan M-W. 2008. Pseudomonas aeruginosa Suppresses Host Immunity by Activating the DAF-2 Insulin-Like Signaling Pathway in Caenorhabditis elegans. PLoS Pathog 4:e1000175.

38. Irazoqui JE, Troemel ER, Feinbaum RL, Luhachack LG, Cezairliyan BO, Ausubel FM. 2010. Distinct Pathogenesis and Host Responses during Infection of C. elegans by P. aeruginosa and S. aureus. PLoS Pathog 6:e1000982.

39. Darnaud M, Dos Santos A, Gonzalez P, Augui S, Lacoste C, Desterke C, De Hertogh G, Valentino E, Braun E, Zheng J, Boisgard R, Neut C, Dubuquoy L, Chiappini F, Samuel D, Lepage P, Guerrieri F, Doré J, Bréchot C, Moniaux N, Faivre J. 2018. Enteric Delivery of Regenerating Family Member 3 alpha Alters the Intestinal Microbiota and Controls Inflammation in Mice With Colitis. Gastroenterology 154:1009–1023.e14.

40. Hanson MA, Grollmus L, Lemaitre B. 2023. Ecology-relevant bacteria drive the evolution of host antimicrobial peptides in *Drosophila*. Science 381:eadg5725.

41. Yu H, Lai H-J, Lin T-W, Chen C-S, Lo SJ. 2015. Loss of DNase II function in the gonad is associated with a higher expression of antimicrobial genes in Caenorhabditis elegans. Biochem J 470:145–154.

42. Yamazaki Y, Akashi R, Banno Y, Endo T, Ezura H, Fukami-Kobayashi K, Inaba K, Isa T, Kamei K, Kasai F, Kobayashi M, Kurata N, Kusaba M, Matuzawa T, Mitani S, Nakamura T, Nakamura Y, Nakatsuji N, Naruse K, Niki H, Nitasaka E, Obata Y, Okamoto H, Okuma M, Sato K, Serikawa T, Shiroishi T, Sugawara H, Urushibara H, Yamamoto M, Yaoita Y, Yoshiki A, Kohara Y. 2010. NBRP databases: databases of biological resources in Japan. Nucleic Acids Res 38:D26–D32.

43. Stiernagle T. 2006. Maintenance of C. elegans. WormBook 10.1895/wormbook.1.101.1.

44. Chen S, Zhou Y, Chen Y, Gu J. 2018. fastp: an ultra-fast all-in-one FASTQ preprocessor. Bioinformatics 34:i884–i890.

45. Bray NL, Pimentel H, Melsted P, Pachter L. 2016. Near-optimal probabilistic RNA-seq quantification. 5. Nat Biotechnol 34:525–527.

46. Pimentel H, Bray NL, Puente S, Melsted P, Pachter L. 2017. Differential analysis of RNA-seq incorporating quantification uncertainty. 7. Nat Methods 14:687–690.

47. Higgins DP, Weisman CM, Lui DS, D’Agostino FA, Walker AK. 2022. Defining characteristics and conservation of poorly annotated genes in Caenorhabditis elegans using WormCat 2.0. Genetics 221:iyac085.

48. Trang K, Bodkhe R, Shapira M. 2022. Compost Microcosms as Microbially Diverse, Natural-like Environments for Microbiome Research in Caenorhabditis elegans. JoVE J Vis Exp e64393.

49. Callahan BJ, McMurdie PJ, Rosen MJ, Han AW, Johnson AJA, Holmes SP. 2016. DADA2: High-resolution sample inference from Illumina amplicon data. Nat Methods 13:581–583.

50. Wright ES. 2015. DECIPHER: harnessing local sequence context to improve protein multiple sequence alignment. BMC Bioinformatics 16:322.

51. McMurdie PJ, Holmes S. 2013. phyloseq: An R Package for Reproducible Interactive Analysis and Graphics of Microbiome Census Data. PLOS ONE 8:e61217.

52. Schliep KP. 2011. phangorn: phylogenetic analysis in R. Bioinformatics 27:592– 593.

53. Dixon P. 2003. VEGAN, a package of R functions for community ecology. J Veg Sci 14:927–930.

54. Wilson N, Zhao N, Zhan X, Koh H, Fu W, Chen J, Li H, Wu MC, Plantinga AM. 2021. MiRKAT: kernel machine regression-based global association tests for the microbiome. Bioinformatics 37:1595–1597.

55. Schindelin J, Arganda-Carreras I, Frise E, Kaynig V, Longair M, Pietzsch T, Preibisch S, Rueden C, Saalfeld S, Schmid B, Tinevez J-Y, White DJ, Hartenstein V, Eliceiri K, Tomancak P, Cardona A. 2012. Fiji: an open-source platform for biological-image analysis. 7. Nat Methods 9:676–682.

